# The duration of antibiotic treatment is associated with carriage of toxigenic and atoxigenic strains of *Clostridioides difficile* in dogs

**DOI:** 10.1101/2021.01.12.426335

**Authors:** C. Albuquerque, D. Pagnossin, K Landsgaard, J. Simpson, D. Brown, J.J. Irvine, D. Candlish, A.E. Ridyard., G Douce, C Millins

**Author notes:** Department of Livestock and One Health, Institute of Infection, Veterinary and Ecological Sciences, University of Liverpool (C Millins). Department of Veterinary Pathobiology, College of Veterinary medicine and Biomedical sciences, Texas A&M University, College Station, Texas (K Landsgaard). Joint Corresponding authors.

## Abstract

*Clostridioides difficile* is a leading cause of human antibiotic-associated diarrhoeal disease globally. Zoonotic reservoirs of infection are increasingly suspected to play a role in the emergence of this disease in the community and dogs are considered as one potential source. Here we use a canine case-control study at a referral veterinary hospital in Scotland to assess: i) the risk factors associated with carriage of *C. difficile* by dogs, ii) whether carriage of *C. difficile* is associated with clinical disease in dogs and iii) the similarity of strains isolated from dogs with local human clinical surveillance. The overall prevalence of *C. difficile* carriage in dogs was 18.7% (95% CI 14.8-23.2%, n=61/327) of which 36% (n=22/61) were toxigenic strains. We found risk factors related to prior antibiotic treatment were significantly associated with *C. difficile* carriage by dogs. However, the presence of toxigenic strains of *C. difficile* in a canine faecal sample was not associated with diarrhoeal disease in dogs. Active toxin was infrequently detected in canine faecal samples carrying toxigenic strains (2/11 samples). Both dogs in which active toxin was detected had no clinical evidence of gastrointestinal disease. Among the ten toxigenic ribotypes of *C. difficile* detected in dogs in this study, six of these (012, 014, 020, 026, 078, 106) were ribotypes commonly associated with human clinical disease in Scotland, while atoxigenic isolates largely belonged to 010 and 039 ribotypes. Whilst *C. difficile* does not appear commonly associated with diarrhoeal disease in dogs, antibiotic treatment increases carriage of this bacteria including toxigenic strains commonly found in human clinical disease.

## Introduction

*Clostridioides (formerly Clostridium) difficile* has emerged as a leading cause of antibiotic associated diarrhoeal disease in people globally which is associated with significant morbidity, mortality and healthcare costs (1). In the past this disease was predominately associated with elderly patients treated with antibiotics in healthcare environments, however sentinel surveillance studies have revealed that a substantial proportion of *C. difficile* infections (CDI) are acquired within the community (2–5). Whole genome sequencing has shown that only around a third of hospital cases can be linked to horizontal transmission from symptomatic patients, with the remainder caused by diverse strains of *C. difficile* (6). The source of these infections, and those arising within the community is unknown, and may include asymptomatic human carriers, zoonotic reservoirs, food and the environment (7–9).

Most research on *C. difficile* in animals has focused on production animals and horses with emergence of the 078 ribotype as a significant cause of enteritis in piglets and adult horses occurring around the same time as emergence of CDI in humans (10). The frequent isolation of this organism from the faeces of production animals including pigs, cattle and chickens and food has led to concerns that spread to humans can occur through contamination of the local environment or via the food chain (11–15). Whole genome sequencing of identical strains of *C. difficile* in pig farmers and pigs on the same farm suggests that interspecies transmission is likely, although a common environmental source cannot be excluded (16). In contrast, less is known about the potential of companion animals including dogs, to become colonised with *C. difficile*, develop disease or act as a zoonotic reservoir. The frequency of dog ownership and close living relationships with people justifies evaluation of this species as a potential reservoir host of zoonotic strains of *C. difficile*.

Results from published studies of *C. difficile* carriage by companion animals report prevalence rates in dogs from 0% to 58%, with a lower prevalence in healthy dogs (17–19) and a higher prevalence reported in hospitalised dogs (20) and those visiting human hospitals (21). Similar ribotypes have been identified in both canines and humans suggesting potential for interspecies transmission (22,23). A small number of studies have looked at risk factors for *C. difficile* carriage in dogs. These risk factors may increase individual susceptibility to colonisation such as antibiotic treatment, or be potential sources of infection such as diet and household factors (24–28). Results to date are often contradictory, which may reflect differences in study design and geographic location.

Similarly, existing studies investigating associations between *C. difficile* and disease in dogs have found contrasting results. *C. difficile* in humans is largely a toxin-mediated disease with most pathogenic isolates of *C. difficile* producing one or both major toxins: toxin A (an enterotoxin) and toxin B (a cytotoxin) (29). Atoxigenic and toxigenic strains of *C. difficile* have been detected by several studies in both healthy dogs and those with diarrhoea by bacterial culture and PCR testing for toxin genes (24,30,31). To assess associations between carriage of toxigenic strains and clinical disease, testing for the presence of active toxin in faecal samples from healthy dogs and those with diarrhoea is also needed (17). Some studies which have tested for active toxin have suggested an association between the presence of toxin in faecal samples and diarrhoeal disease in dogs (17,32,33). However, others have not which may in part be explained by differences in methods used to detect active toxin in samples (34).

To assess the potential role of dogs as a zoonotic reservoir of *C. difficile*, and association with clinical diarrhoeal disease in dogs we designed a case-control study of dogs presenting to a veterinary referral practice in Scotland. Our study had the following objectives; i) to assess the risk factors associated with carriage of *C. difficile* by dogs, ii) to test for associations between carriage of *C. difficile* and diarrhoeal disease in dogs and iii) to determine if dogs carry strains of *C. difficile* that are frequently associated with clinical disease in humans.

## Materials and methods

Ethical approval for the study was obtained from the University of Glasgow, School of Veterinary Medicine Ethics and Welfare Committee (Reference number 11a/16). To investigate whether *C. difficile* carriage was associated with disease in dogs we recruited a total of 327 dogs referred from across the west of Scotland to a companion animal veterinary referral hospital. A referral hospital was chosen for the study due to the large geographic catchment area, ability to include dogs with a range of potential risk factors for carriage of *C. difficile* and the capacity to investigate the association of *C. difficile* carriage with diarrhoeal disease. A fresh faecal sample collected by the owner on the day of admission to the hospital or a sample from the first stool passed within 48 hours of admission was used to evaluate *C. difficile* colonisation of dogs within the community (24). Dogs were referred for a range of specialist medical or surgical reasons to the University of Glasgow School of Veterinary Medicine Small Animal Hospital (hereafter described as the referral hospital). Dogs referred for treatment of either acute or chronic diarrhoea (n=101) or for non-gastrointestinal reasons (n=226) were recruited to the study between June 2016 and October 2019.

### Assessment of risk factors for C. difficile carriage by dogs

A questionnaire designed to assess potential risk factors for the carriage of *C. difficile* by dogs was completed by a subset of owners (n=200) recruited to the study. All owners presenting to the referral hospital with their dog between June to December 2016 were invited to complete the survey unless the dog was critically ill. The survey was designed to provide information on potential risk factors for increased susceptibility for *C. difficile* carriage and to identify potential sources of infection. We requested information on the diet, breed, sex and age of the dog and information on the household, including co-habiting pets, elderly people or infants. Clinical information on antibiotic, antacid, immunosuppressive treatment and the number of visits to and days as an inpatient at a veterinary hospital within the three months prior to admission to the referral hospital was obtained by examining the clinical records from the referring practice.

### Detection and strain typing of C. difficile from canine faecal samples

Following collection, canine faecal samples were placed in anaerobic jars and processed within 72 hours. One gram of each faecal sample was emulsified in 2mls of ethanol and incubated at room temperature for 30 minutes to kill vegetative cells and induce sporulation of *C. difficile*. Two hundred microlitres of faecal suspension was plated onto Brazier’s taurocholate cycloserine cefoxitin agar (TCCA) supplemented with 5% defibrinated horse blood and egg white emulsion. Each sample was cultured in duplicate and incubated for a maximum of 7 days in an anaerobic chamber (Don Whitley, UK). Colonies showing typical *C. difficile* colony morphology and which appeared black when subcultured onto Biomerieux chromID^®^ *C. difficile* agar were selected. Clones were identified as *C. difficile* by amplification and analysis of the 16S (V4) ribosomal RNA sequence (Supporting information, Table 1). Chromosomal DNA was prepared from each isolate using DNeasy Blood and Tissue kits, (Qiagen, Hilden, Germany), following the manufacturer’s instructions. Amplified PCR products were purified using the Qiaquick PCR purification kit, and sequenced by Source Bioscience (Livingston, UK). The bacterial species for each isolate was determined by subjecting each sequenced and trimmed PCR product to BLAST analysis using the National Centre for Biotechnology (NCBI) Nucleotide database.

**Table 1:** Univariable analysis of risk factors potentially associated with the carriage of *C. difficile* in dogs presented to the University of Glasgow, Small Animal Hospital referral hospital.

| Explanatory Variable | Sample | Factor level | Odds ratio (95% CI) | p value* |
| --- | --- | --- | --- | --- |
| Sex and neutering status | 5/29 | Female | 1.0 (reference) | 0.92 |
|  | 8/55 | Female (N) | 0.82 (0.24-2.95) |  |
|  | 10/51 | Male | 1.17 (0.37-4.13) |  |
|  | 11/61 | Male (N) | 0.98 (0.32-3.38) |  |
| Age (months) | 200 | NA | 1.01 (1.002-1.02) | 0.016 |
| Raw meat in diet | 32/182 | No | 1.0 (reference) | 0.46 |
|  | 2/18 | Yes | 0.59 (0.09-2.20) |  |
| Cooked meat in diet | 21/119 | No | 1.0 (reference) | 0.77 |
|  | 13/81 | Yes | 0.89 (0.41-1.88) |  |
| Other pets in household | 21/98 | No | 1.0 (reference) | 0.10 |
|  | 13/102 | Yes | 0.54 (0.25-1.13) |  |
| Number of people in household | 18/106 | Two or less | 1.0 (reference) | 0.94 |
|  | 16/92 | More than two | 1.03 (0.49-2.16) |  |
| Infant (< 2 years) or person > 65 years in household | 29/174 | No | 1.0 (reference) | 0.75 |
|  | 5/26 | Yes | 1.19 (0.37-3.21) |  |
| Antibiotic treatment last 3 months | 10/99 | No | 1.0 (reference) | 0.009 |
|  | 24/101 | Yes | 2.77 (1.28-6.42) |  |
| Length of antibiotic treatment (days) | 200 | NA | 1.08 (1.04-1.11) | <0.001 |
| Treatment with more than 1 antibiotic | 200 | No antibiotic | 1.0 (reference) | 0.024 |
|  |  | One antibiotic | 2.2 (0.89-5.58) |  |
|  |  | >1 antibiotic | 3.52 (1.38-9.16) |  |
| Immunosuppressive treatment | 30/182 | No | 1.0 (reference) | 0.55 |
|  | 4/18 | Yes | 1.45 (0.39-4.37) |  |
| Antacid treatment | 23/153 | No | 1.0 (reference) | 0.19 |
|  | 11/47 | Yes | 1.73 (0.75-3.81) |  |
| Overnight stay in veterinary hospital | 20/131 | No | 1.0 (reference) | 0.33 |
|  | 14/67 | Yes | 1.47 (0.68-3.11) |  |
| Visits to veterinary hospital | 6/48 | One or less | 1.0 (reference) | 0.54 |
|  | 20/104 | 2-5 | 1.71 (0.67-4.96) |  |
|  | 8/48 | >5 | 1.40 (0.45-4.59) |  |

\*p value from likelihood ratio tests compared to a null model

All isolates were ribotyped at the Scottish Microbiology Reference Laboratory (Glasgow) (35). Ribotype patterns were assigned following analysis using BioNumerics v7.6 software (Applied Maths, Sint-Maartens-Latem, Belgium). Patterns were compared with libraries using the Pearson Correlation Coefficient of similarity with 1% optimisation.

### Detection of toxin by PCR assays for tcd A and tcd B and cytotoxicity assay

All isolates of *C. difficile* were tested for the presence of fragments of the *tcd*A and *tcd*B genes by PCR amplification with primers designed using the annotated *C. difficile* 630 genome (Supporting information, Table 1). Amplification was shown to be specific by inclusion of DNA from the epidemic, toxin-producing strain *C. difficile* R20291 and from a PaLoc negative strain, 1342.

A subset of 116 faecal samples collected between 2018-19, were tested for the presence of active toxin using a cytotoxicity assay. Of these samples, 22 were from dogs presenting with diarrhoea (8 with acute diarrhoea and 14 with chronic diarrhoea) and 94 samples were from control dogs without diarrhoea. To detect the presence of toxin, aliquots of fresh faeces within 48 hours of collection, were emulsified in 2ml of PBS and solid material removed by centrifugation. The supernatant was filtered through a 0.2um membrane filter and the resultant material was serially diluted in PBS and added to a prepared monolayer of Vero cells. To confirm specificity of cytotoxicity, a second set of samples, prepared in parallel, were treated with *Clostridium sordellii* antitoxin (NIBSC, 20 IU/ml). This antitoxin cross-reacts with *C. difficile* toxin and has been used to confirm the presence of *C. difficile* toxin activity in human samples (36). Treated cells were then incubated for 18-24 hours at 37°C with 5% CO_2_, before cells were fixed with 1% formalin for 30 minutes and stained using Giemsa stain (SIGMA-ALDRICH®, 6% diluted) for 1 hour. Cell rounding, which is associated with *C. difficile* toxin presence, was assessed microscopically and samples were considered positive if cell rounding was observed that was neutralized by addition of the *C. sordellii* antitoxin (36).

### Statistical analysis

All statistical analyses were carried out in R version 4.0.2 (R Development Core Team, Vienna, Austria using the package *lme4* (37) for analyses. Collinearity was tested for using the variance inflation factor in the *car* package in ‘R’ (38). Prevalence was calculated using the prop.test function in ‘R’.

### Assessment of risk factors for carriage of C. difficile

*C. difficile* carriage in dogs (present or absent) was modelled in a binomial general linear model (GLM) with a logit link as a function of each of the following potential risk factors listed in Table 1. Risk factors with a p value of < 0.10 based on univariable analysis were included in a multivariable general linear model of *C. difficile* carriage (present or absent) with binomially distributed errors and a logit link. Starting from the maximum global model, stepwise backwards model selection was carried out using likelihood ratio tests.

### Testing for an association between C. difficile carriage and diarrhoeal disease in dogs

To test whether the carriage of toxigenic strains of *C. difficile* was associated with diarrhoea, the presence or absence of diarrhoea in each dog (n=327) was modelled using a GLM with binomially distributed errors and a logit link, as a function of *C. difficile* carriage, and separately as a function of carriage of a toxigenic strain of *C. difficile* (based on the PCR presence of one or more toxin genes).

### Comparison of strains detected in dogs with clinical surveillance for C. difficile in humans

To assess the potential for shared strains between dogs and humans, ribotypes detected from dogs in this study were compared to ribotypes recorded from human surveillance of *C. difficile* cases by the National Microbiology Reference laboratory in Scotland between 2015 and 2018.

## Results

### Prevalence and strain diversity of C. difficile in canine faecal samples

The overall prevalence of *C. difficile* from canine faecal samples in this study was 18.7% (95% 14.8 – 23.2%, n = 61/327). The majority of isolates were atoxigenic strains (63.4% n = 39/61) while 22/61 were toxigenic strains with the PCR presence of either or both *tcd*A and *tcd*B genes. A total of 13 different ribotypes were detected in the canine samples, and 10 of these ribotypes included toxigenic isolates (Figure 1, Supporting information Table 2).

**Table 2:** Results from the final selected multivariable general linear model to explain carriage of *C. difficile* by dogs.

| Model description | Fixed effects | Mean<br>(Estd) | SE | Odds ratio<br>(95% CI) | p value* |
| --- | --- | --- | --- | --- | --- |
| *C. difficile* carriage<br>(presence/absence) | Intercept | -2.1 | 0.25 | NA | NA |
|  | Length of antibiotic treatment<br>(days) | 0.047 | 0.012 | 1.08 (1.04-1.11) | <0.001 |

\*p value from LRT

**Figure 1:**
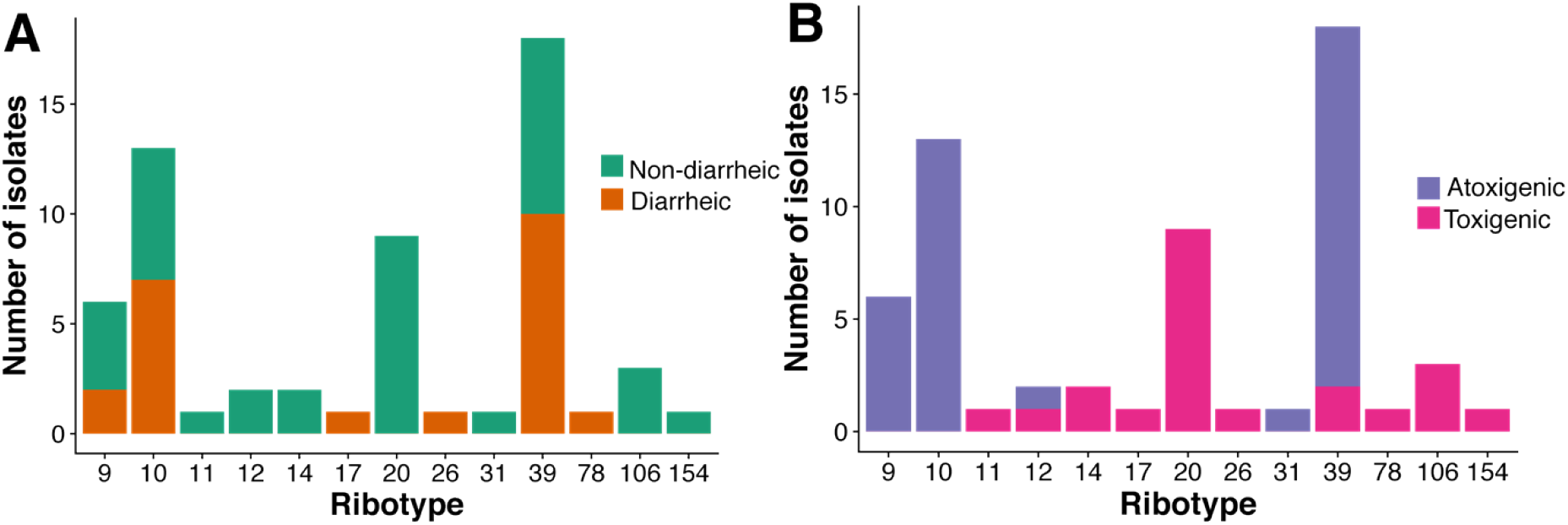
Ribotypes of *C. difficile* from dogs in this study, shown according to whether they were isolated from dogs with or without diarrhoea (A) and whether strains were classed as toxigenic or atoxigenic strains (B) based on PCR testing *tcd*A and *tcd*B genes.

### Cytotoxicity Assay

A total of 116 faecal samples were tested for active toxin, and two samples tested positive. Both faecal samples with active toxin present were from dogs with no clinical evidence of diarrhoea. *C. difficile* was subsequently cultured from 26 of the 116 faecal samples tested for active toxin (21 dogs without diarrhoea, and 5 dogs with diarrhoea) and 11 of these isolates tested PCR positive for either or both *C. difficile* toxin genes (*tcd*A and *tcd*B). This included both samples which tested positive with the cytotoxicity assay. Ribotype analysis revealed these strains to be 020 and 106 types respectively (Supporting information, Table 2).

### Risk factors for carriage of C. difficile by dogs

Results of univariable analysis of risk factors and carriage of *C. difficile* are shown in Table 1. In a multivariable model, dogs were more likely to carry *C. difficile* with an increasing length of treatment on antibiotics, for each day of antibiotic treatment (OR = 1.08, 95% C.I. 1.04 − 1.11), p<0.001 (Table 2). Other explanatory variables including age (months), treatment with multiple antibiotics and the presence of multiple pets in the household were not maintained in the final model.

### Testing for an association between C. difficile carriage and diarrhoeal disease in dogs

Neither carriage of *C. difficile* (OR = 1.33 95% CI = 0.74 − 2.38, p = 0.34), or the presence of toxigenic strains of *C. difficile* in a faecal sample (OR = 0.60, 95% CI = 0.19-1.55, p = 0.31) was associated with diarrhoea in dogs. Toxigenic strains of *C. difficile* were detected both in dogs with diarrhoea and in dogs with no evidence of gastrointestinal disease (Figure 1).

### Comparison of strains detected in dogs with clinical surveillance for C. difficile in humans

The most commonly detected ribotypes from human clinical surveillance during the time period of the study are shown in Supporting information Table 3. Six of these ribotypes, 012, 014, 020, 026, 078, 106 were isolated from dogs in this study.

## Discussion

This study has found that *C. difficile* can be frequently isolated from diarrheic and non-diarrheic canine faecal samples, and carriage of toxigenic strains by dogs is not associated with diarrhoeal disease. As in humans and other species, antibiotic treatment was significantly associated with the carriage of *C. difficile* by dogs. Several toxigenic ribotypes detected in dogs in this study are among the most frequently reported ribotypes from clinical surveillance of people in the same locality over the period of the study.

The overall prevalence of *C. difficile* carriage in dogs in this study was 18.7%, similar to previous studies from referral hospitals in other countries which reported rates of 18.4% and 13.7% (20,26). Prevalence rates in healthy dogs presenting to primary care veterinary clinics and those living in shelters are reported to be lower, ranging from 0% to 6.1% (19,30). Several factors could contribute to these apparent differences, including geographic area, laboratory methods and age and clinical history of dogs recruited to these studies. In our study toxigenic isolates accounted for 36% of positive cultures (n=22/61); previous studies have reported a prevalence of toxigenic isolates of 36.8% to 69% (20,22,24,27). The most commonly isolated ribotypes in our study cohort were ribotypes 039 and 010, followed by 020, a toxigenic ribotype that was recovered most frequently from non-diarrheic dogs (Figure 1). In contrast to studies of production animals and horses where the 078 ribotype often dominates, no dominate strain appears to be present across canine studies (19,23,25,28,39,40). Other European studies have reported the 010 ribotype as being among the most common strains isolated from dogs including those from Germany, Holland, Spain and Italy (19,23,28,39,40). In these studies, ribotypes 014 and 020 were also frequently isolated.

We found risk factors related to antibiotic treatment within the previous three months were significantly associated with carriage of *C. difficile* by dogs. The length of antibiotic treatment was the only factor supported in a multivariable model, with a seven-day course of treatment predicted to increase the risk of carriage by 1.67 times (95% CI 1.30-2.13, p <0.001). Some previous studies in dogs have found an association between previous antibiotic administration and *C. difficile* carriage (26,28,41) whereas other studies did not (20,24,27). As in humans and horses the mechanism underpinning the positive relationship between antibiotic treatment and *C. difficile* carriage is likely due to loss of microbiome diversity within the gut (42). As a result, *C. difficile* is able to germinate and rapidly multiply in the available niche. Age was also positively associated with carriage of *C. difficile* on univariable analysis with a slightly increased risk of carriage per year OR =1.13 (95% CI=1.02-1.25, p=0.016), as reported by other studies (20,27). This positive relationship could suggest an extended duration of colonisation, though no longitudinal studies of carriage in dogs have been carried out to date. Alternatively, there could be increased host susceptibility with age. We were unable to identify a potential source of *C. difficile* colonisation of dogs from our questionnaire survey. Previous studies have found that dogs living with an immunocompromised person or contact with a person with diarrhoea can increase the risk of colonisation in dogs while feeding a dry food diet reduces risk (18,28,43,44). Although comparison of the results of published studies in dogs is limited by difference in study design and geographic area, a potential explanation for variation in ribotypes detected in dogs could be that carriage is driven mainly by host susceptibility. If this hypothesis is true, the strains isolated from dogs may be reflective of those which they are exposed to on a daily basis in food and the environment (9,14).

The significance of *C. difficile* as cause of disease in dogs is unclear, since toxigenic strains can be isolated from healthy, as well as diarrheic dogs (24,30). Our study was in agreement with others which did not find an association between carriage of toxigenic strains of *C. difficile* and diarrhoea (24,33). Using a cytotoxicity assay, we found that production of active toxin in dogs was uncommon when toxigenic strains were isolated. This was also found by another recent study (45). In our study only two of eleven samples which carried toxigenic strains tested positive for active toxin and both of these samples were from non-diarrheic dogs. Some previous studies which indicated a relationship between the presence of active toxin and *C. difficile* associated disease in dogs may have been affected by low sensitivity and specificity of ELISA’s used to detect toxin in dogs (34). We were limited in our study cohort in evaluating associations between diarrhoea and carriage of toxigenic strains of *C. difficile* due to the relatively low numbers of diarrheic samples carrying toxigenic strains of *C. difficile* (n=5). One of these samples was tested with the cytotoxicity assay and tested negative. Due to time and logistical constraints we were only able to implement the cytotoxicity assay for part of the study period. Although *C. difficile* may still be a potential cause of diarrhoea in dogs, our results suggest that the frequency of disease is likely to be low. The availability of reliable tests for active toxin which are suitable for use in a diagnostic laboratory setting is likely to limit clinical investigations into the significance of toxigenic isolates. Cytotoxicity assays are labour intensive and unlikely to be widely available.

There is evidence that the epidemiology of CDI in humans is changing, with increasing numbers of cases reported from patients residing within the community and attribution of the source of infections in most of these cases is unknown (7,46). This study, in agreement with other recent studies shows that ribotypes associated within human clinical disease can be carried asymptomatically with the canine gut. Six of the ten toxigenic ribotypes of *C. difficile* detected in dogs in this study (012, 014, 020, 026, 078, 106) are also some of the most common isolates detected by human clinical surveillance in Scotland from 2015-2018 (Appendix Table 3) (47–49). A subset of these ribotypes (014, 020 and 078) are amongst the most prevalent causes of *C. difficile*-associated diarrhoea in Europe (50,51). Results from this and other companion animal studies demonstrating shared ribotypes amongst dogs and humans suggest that dogs could contribute to a reservoir for human infections, either directly or by contaminating the environment. Understanding the potential significance of carriage of toxigenic strains of *C. difficile* by companion animals to human community CDI is likely to be challenging and will require integrated molecular epidemiology studies of community CDI with investigation of food, environment and potential zoonotic sources.

### Conclusions

We have found that prevalence of *C. difficile* carriage in dogs presenting to a referral hospital in Scotland is relatively common, and an increasing length of antibiotic therapy is associated with a higher risk of *C. difficile* carriage. The findings of this study and others suggest that *C. difficile* is not commonly associated with diarrhoeal disease in dogs. Dogs carried several toxigenic strains associated with human clinical disease and could potentially act as a source of infection for humans, or spore accumulation within the environment.

## Acknowledgements

This research was supported by a Petplan Charitable Trust pump primer award and the University of Glasgow Small Animal Fund. The authors are grateful for support of staff and technicians at the University of Glasgow Small Animal Hospital with the recruitment of animals and collection of samples for the study and for advice on the statistical analysis from Paul Johnson. KL was supported by a BBSRC STARS scholarship. JS was supported by a vacation scholarship from The Carnegie Trust.

## Contributions

GRD and CM conceived the study with advice from AR, CA and DP collected faecal samples and clinical information, DP performed cytotoxin testing, DP, KL, JI, DC, GRD undertook culture, DNA extraction, toxin and 16S PCR analysis, DB evaluated isolate ribotype, CM carried out the data analysis, CM and GRD produced the first draft of the manuscript, all authors read and approved the final manuscript.

## Supporting Information

**Table 1:**
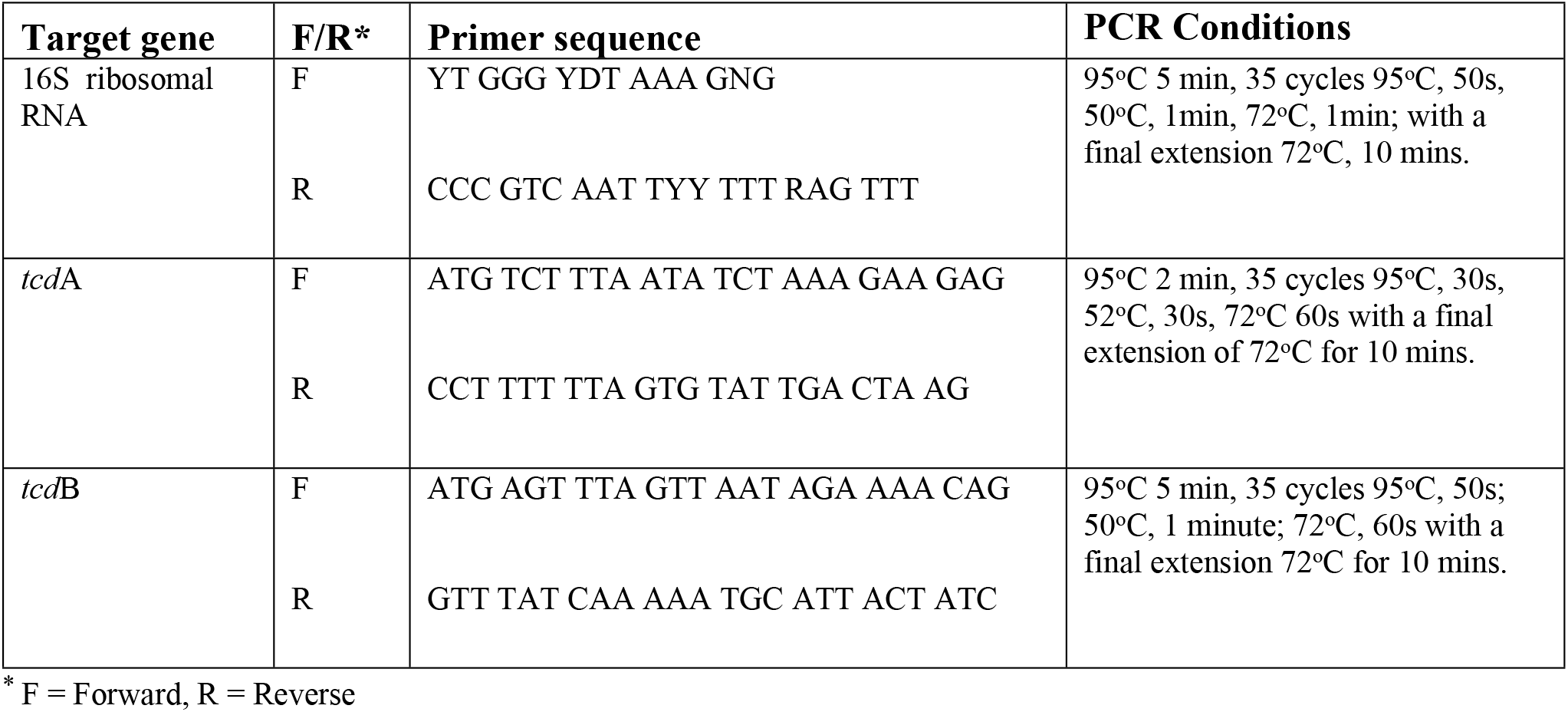
Primers and amplification conditions for 16S and (*tcd*A and *tcd*B) PCR.

**Table 2:** Table to show *C. difficile* isolates from dogs referred for investigation of diarrhoea, or for a non-gastrointestinal complaint. Ribotype and results of *tcd*A and *tcd*B PCR testing and cytotoxicity testing for active toxin presence are shown.

| Clinical presentation | Ribotype | *tcdA* | *tcdB* | Cytotoxicity assay |
| --- | --- | --- | --- | --- |
| Diarrhoea | 009 | 0 | 0 | 0 |
|  | 010 | 0 | 0 | 0 |
|  | 039 | 0 | 0 | 0 |
|  | 039 | 0 | 0 | NA |
|  | 026 | 1 | 1 | 0 |
|  | 039 | 0 | 0 | NA |
|  | 078 | 1 | 1 | NA |
|  | 010 | 0 | 0 | NA |
|  | 010 | 0 | 0 | NA |
|  | 010 | 0 | 0 | NA |
|  | 009 | 0 | 0 | NA |
|  | 010 | 0 | 0 | NA |
|  | 010 | 0 | 0 | 0 |
|  | 017 | 0 | 1 | NA |
|  | 039 | 0 | 0 | NA |
|  | 039 | 1 | 1 | NA |
|  | 039 | 0 | 0 | NA |
|  | 039 | 0 | 0 | NA |
|  | 039 | 0 | 0 | NA |
|  | 039 | 0 | 0 | NA |
|  | 039 | 1 | 1 | NA |
|  | 010 | 0 | 0 | NA |
| No<br>diarrhoea | 011 | 1 | 1 | 0 |
|  | 009 | 0 | 0 | 0 |
|  | 039 | 0 | 0 | 0 |
|  | 020 | 1 | 1 | 0 |
|  | 009 | 0 | 0 | 0 |
|  | 106 | 1 | 1 | 1 |
|  | 010 | 0 | 0 | 0 |
|  | 154 | 1 | 1 | 0 |
|  | 039 | 0 | 0 | 0 |
|  | NA | 0 | 0 | NA |
|  | 039 | 0 | 0 | NA |
|  | 020 | 1 | 1 | NA |
|  | NA | NA | NA | NA |
|  | 039 | 0 | 0 | NA |
|  | 020 | 1 | 1 | NA |
|  | 020 | 1 | 1 | NA |
|  | 020 | 1 | 1 | NA |
|  | 009 | 0 | 0 | 0 |
|  | 031 | 0 | 0 | 0 |
|  | 106 | 1 | 1 | 0 |
|  | 012 | 1 | 1 | 0 |
|  | 010 | 0 | 0 | 0 |
|  | 020 | 1 | 1 | 0 |
|  | 039 | 0 | 0 | 0 |
|  | 009 | 0 | 0 | 0 |
|  | 020 | 1 | 1 | 1 |
|  | 020 | 0 | 1 | 0 |
|  | 039 | 0 | 0 | 0 |
|  | 014 | 1 | 1 | 0 |
|  | 010 | 0 | 0 | NA |
|  | 010 | 0 | 0 | NA |
|  | 039 | 0 | 0 | NA |
|  | 106 | 1 | 1 | NA |
|  | 014 | 1 | 1 | NA |
|  | 010 | 0 | 0 | NA |
|  | 012 | 0 | 0 | NA |
|  | 010 | 0 | 0 | NA |
|  | 039 | 0 | 0 | NA |
|  | 020 | 1 | 1 | NA |

**Table 3:** Human clinical surveillance for *Clostridium difficile* in Scotland. Data from Health Protection Scotland Annual Reports 2016-2018 (1-3).

| Ribotype | 2015 |  | 2016 |  | 2017 |  | 2018 |  |
| --- | --- | --- | --- | --- | --- | --- | --- | --- |
|  | Number | % | Number | % | Number | % | Number | % |
| **1** | 015 | 3.9 | 10 | 3.2 | NA | NA | NA | NA |
| **2** | 050 | 13.2 | 32 | 10.4 | 36 | 10.4 | 45 | 12.9 |
| **5** | 033 | 8.7 | 23 | 7.4 | 32 | 9.2 | 26 | 7.4 |
| **12** | 015 | 3.9 | 1 | 0.3 | NA | NA | NA | NA |
| **14** | 032 | 8.4 | 33 | 10.7 | 32 | 9.2 | 31 | 8.9 |
| **15** | 035 | 9.2 | 22 | 7.1 | 32 | 9.2 | 34 | 9.7 |
| **20** | 025 | 6.6 | 22 | 7.1 | 23 | 6.6 | 26 | 7.4 |
| **23** | 015 | 3.9 | 19 | 6.1 | 15 | 4.3 | 28 | 8 |
| **26** | 013 | 3.4 | 5 | 1.6 | NA | NA | NA | NA |
| **27** | NA | NA | NA | NA | 1 | 0.3 | 2 | 0.6 |
| **78** | 040 | 10.5 | 25 | 8.1 | 34 | 9.8 | 41 | 11.7 |
| **106** | NA | NA | NA | NA | 13 | 3.8 | 8 | 2.3 |
| **Others** | 107 | 28.2 | 117 | 37.9 | 128 | 37 | 108 | 30.9 |

